# Aberrant axon initial segment is an indicator for task-independent neural activity in autism model mice

**DOI:** 10.1101/2024.02.15.580571

**Authors:** Yoshinori Otani, Kohei Koga, Ryo Kawabata, Xinglang Liu, Hisao Miyajima, Toru Takumi, Masashi Fujitani

**Affiliations:** Department of Anatomy and Neuroscience, Faculty of Medicine, Shimane University, Izumo, Japan; Department of Neurophysiology, Faculty of Medicine, Hyogo Medical University, Nishinomiya, Hyogo, Japan; Gladstone Institute of Neurological Disease, San Francisco, USA; Department of Physiology and Cell Biology, Kobe University School of Medicine, Chuo, Kobe, Japan

## Abstract

Autism spectrum disorder (ASD) is a developmental disorder characterized by impairments in several communicative behaviors. Identification of biomarkers for ASD based on task-independent neural activity has been elusive. The axonal initial segment (AIS), located at the proximal part of the axon, is crucial for initiating action potentials (APs) and adaptable through activity-dependent AIS plasticity. In our study, we discovered decrease in AIS length in layer V pyramidal neurons (PyNs) of the prelimbic cortex (PrL) and increase in layer II/III PyNs in the somatosensory cortex of duplicated human *15q11-13* region (*15q dup*) ASD model mice. Electrophysiological studies using whole-cell patch-clamp recordings in the PrL revealed diminished ability to generate APs. We discovered lack of AIS plasticity in 15*q dup* mice under conditions of elevated potassium chloride. Retrograde tracing demonstrated that AIS shortening depends on the projection targets. These results indicate that activity-dependent aberrant AIS structures can serve as indicators of altered task-independent neural activity in ASD mouse models.

## INTRODUCTION

The axon initial segment (AIS) is located proximal to the axon and distinguished by the high density of the AIS anchoring protein, Ankyrin-G, and ion channels, which play a pivotal role in initiating action potentials (APs)^1, 2, 3, 4^. AIS is a critical regulator of neuronal excitability, serving as an initiator of AP. Alterations in its structural features, such as length and position, have been shown to influence neuronal function^1, 2, 3, 4^. This concept, known as activity-dependent AIS plasticity^5, 6^, has been demonstrated in earlier *in vivo* studies, utilizing sensory deprivation paradigms such as tactile^7^, auditory^8, 9^, visual^10, 11^ and olfactory sensations^12^. Beyond the physiological aspects, an expanding volume of research has linked abnormalities in the AIS to a spectrum of neurological disorders, including neuropathic pain^13^, Alzheimer’s disease^14^, stroke^15^, brain abnormality in type 2 diabetes^16^, attention-deficit hyperactivity disorder^17^ and peripheral nerve injury^18^. Specifically, in animal models of Fragile X syndrome, an increase in cellular excitability with changes in AIS plasticity has been found^19^. In animal models of Angelman syndrome, Ankyrin-G expression is altered, indicating that changes in AIS length might be attributable to abnormal excitability in neurons, as well as to biochemical alterations of AIS-specific proteins such as Ankyrin-G^20^. Moreover, frontotemporal dementia-causing *Tau* mutations have been found to abolish AIS plasticity and impair neuronal activity through AIS cytoskeletal changes effected by end-binding protein 3 (EB3)^21^. These results suggest that AIS changes depend on the neuronal subtype, presence of pathological conditions, and altered neural circuit activity.

Autism spectrum disorder (ASD) is a complex developmental disorder characterized by impairments in communication and reciprocal social interactions, restrictive repetitive behaviors or interests, sensory abnormalities, and varying levels of intellectual disability^22, 23^. Among the genetic factors implicated in ASD, copy number variations (CNVs) are frequently observed chromosomal abnormalities and represent significant risk factors for the disorder^24^. Specifically, CNVs within the *15q11-13* locus have been identified as genomic variants associated with ASD^25^. Moreover, mouse models duplicating the human *15q11-13* region (*15q dup*) have been developed^26^, and these mice exhibit ASD-like symptoms^10^, increased activity in the somatosensory cortex and decreased serotonergic neuron activity^27^.

ASD is characterized by complex dysfunction spanning multiple brain regions, cell types, and neural circuits^28, 29^. To understand how brain-wide activity patterns contribute to and manifest ASD-like behaviors, previous studies have elucidated global and neural circuit dysfunctions in *15q dup* ASD mice models using awake functional magnetic resonance imaging^30^ and real-time imaging^31^. For neural circuit mapping based on task-dependent neural activity, researchers have developed methods to capture neural activity snapshots via immediate early genes such as *Fos* or *Arc*, monitor intracellular Ca^2+^ concentration, and record synaptic events^32, 33^. However, identification of pathological biomarkers based on task-independent neural activity remains elusive. Projection specificity throughout the entirety of the brain is particularly challenging^34^. In this study, we aimed to identify functionally aberrant AIS structures that can serve as valuable biomarkers of task-independent neural activity in an ASD mouse model.

## RESULTS

### Altered AIS length of pyramidal neurons in various brain regions of *15q dup* mice

We first established a precise quantification method to measure AIS length *in vivo*, as shown in Figs. 1 and Supplementary Fig. 1. We visualized cortical pyramidal neurons (PyNs) through the immunohistochemical detection of Ankyrin-G as an AIS marker, combined with layer-specific markers and a neural cell body marker (fluorescent Nissl), in order to accurately identify the location and length of the AIS. Subsequently, we captured three-dimensional neuronal morphology and quantified the AIS length. We selected the following cortical areas to focus on in the *15q dup* mouse: the prelimbic cortex (PrL) and infralimbic cortex (IL) in the medial prefrontal cortex (mPFC), somatosensory cortex (SC), and motor cortex (MC) (Fig. 1a). As demonstrated in Figs. 1b and c, we quantified the AIS length in layers II/III and V PyNs of wild-type (WT) and *15q dup* mice using Ankyrin-G as an AIS marker, along with cortical layer-specific markers, without assessing any behavioral tasks. We observed that the AIS length varied by cell layer and cortical location, with an increase only in layer II/III PyNs of the SC (layer II/III PrL, WT: 32.69 ± 1.103 μm, *15q dup*: 31.43 ± 0.8788 μm; IL, WT: 32.20 ± 1.399 μm, *15q dup*: 31.82 ± 1.318 μm ; SC, WT: 26.47± 0.4864 μm, *15q dup*: 27.68 ± 0.2659 μm; MC, WT: 28.61± 0.8776 μm, *15q dup*: 28.51 ± 0.5293 μm), and a decrease in layer V PyNs of the PrL, compared to WT mice (layer V PrL, WT: 26.62± 0.4889 μm, *15q dup*: 24.49 ± 0.6513 μm; IL, WT: 26.86. ± 0.7923 μm, *15q dup*: 27.37 ± 1.108 μm ; SC, WT: 26.82 ± 0.5559 μm, *15q dup*: 26.23 ± 0.5137 μm; MC, WT: 27.81 ± 0.5191 μm, *15q dup*: 28.55 ± 0.8460 μm), as shown in Figs. 1b, c and Supplementary Fig. 1. As shown in Fig. 1c, the cumulative frequency plots of individual AIS lengths in WT and *15q* mice showed a leftward shift in layer V of the PrL and rightward shift in layers II/III of the SC, and the distribution was comparable with that of controls.

**Fig. 1.**
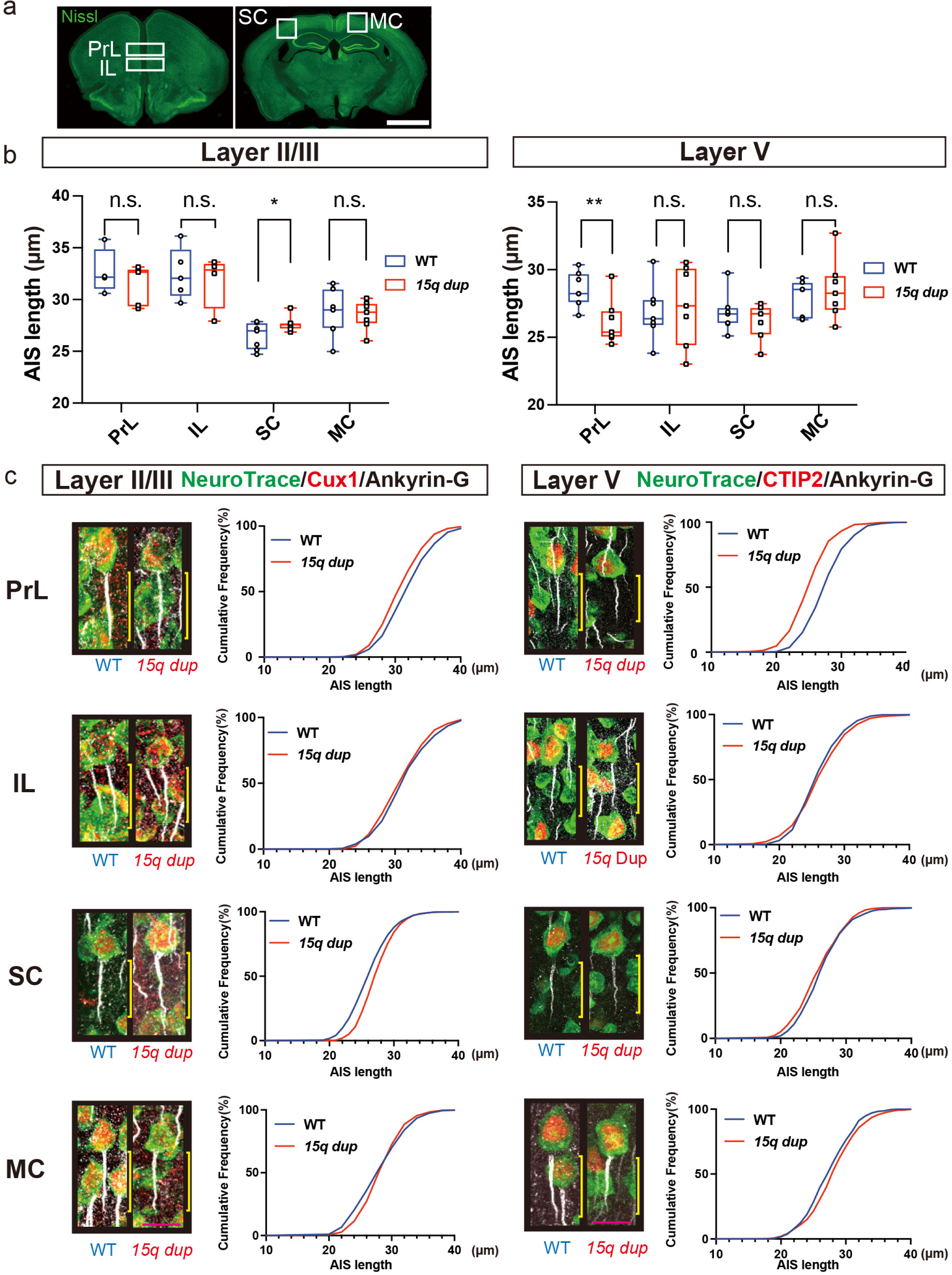
Alteration of the axon initial segment (AIS) length in the layer II/III pyramidal neurons (PyNs) of the somatosensory cortex (SC) and layer V PyNs of the prelimbic cortex (PrL) of duplicated human *15q11-13* region (*15q dup*) autism spectrum disorder (ASD) model mouse. **a)** Representative images of coronal sections of mouse brain stained with fluorescent Nissl, NeuroTrace® (green). White squares indicate areas where the AIS length was measured. Scale bar = 100 µm. **b)** Analysis of AIS length in layer II/III (left) and layer V PyNs (right) in the PrL, IL, SC, and motor cortex (MC) of wild-type (WT) and *15q dup* mouse brains. Values are presented as medians with 25–75% intervals, and error bars show the minimum to maximum values for all data points. n.s., not significant; *P=0.0487, ** P=0.0113, analyzed using two-tailed unpaired Student’s *t-*test (n = 7). Scale bar = 10 µm. **c)** Representative confocal images of PyNs and the cumulative frequency of AIS length in layers II/III (left) and V (right) of WT and *15q dup* mouse. Immunohistochemical images of NeuroTrace®(green), Cux1(red), and Ankyrin-G(white) in the layer II/III PyNs of left panels, and images of NeuroTrace®(green), CTIP2(red), and Ankyrin-G(white) in the layer V PyNs of right panels. Yellow bars indicate the AIS length for each neuron. Scale bar = 10 µm.

### AIS shortening is due to abnormal circuit activity

Next, we investigated whether alteration in AIS length was associated with a misdistribution between voltage-gated sodium (NaV) channels and Ankyrin-G in layer V PrL and layer II/III SC of *15q dup* mice. As depicted in Figs. 2a,b, Pan-NaV channels co-localized with Ankyrin-G, and no significant difference in length was found between clusters of Pan-NaV and those of Ankyrin-G, although the AIS length decreased in the layer V PyNs of the PrL and increased in layer II/III of the SC in the *15q dup* mice, as shown in Fig. 1 (PrL layer V, WT: Ankyrin-G: 27.74 ± 0.4498 μm; Pan-NaV channels: 28.71 ± 0.5280 μm, *15q dup*: Ankyrin-G: 24.33 ± 0.3327 μm; Pan-NaV channels: 24.81 ± 0.4657 μm. SC layer II/III, WT: Ankyrin-G: 25.03 ± 0.6922 μm; Pan-NaV channels: 26.02 ± 0.5518 μm, *15q dup*: Ankyrin-G: 27.26 ± 0.6288 μm; Pan-NaV channels: 27.93 ± 0.4273 μm). Also, no significant difference in length were found in the layers II/III PyNs of the PrL, IL and MC (Supplementary Fig. 2) and, layers V PyNs of the IL,SC and MC (Supplementary Fig. 3).

**Fig. 2.**
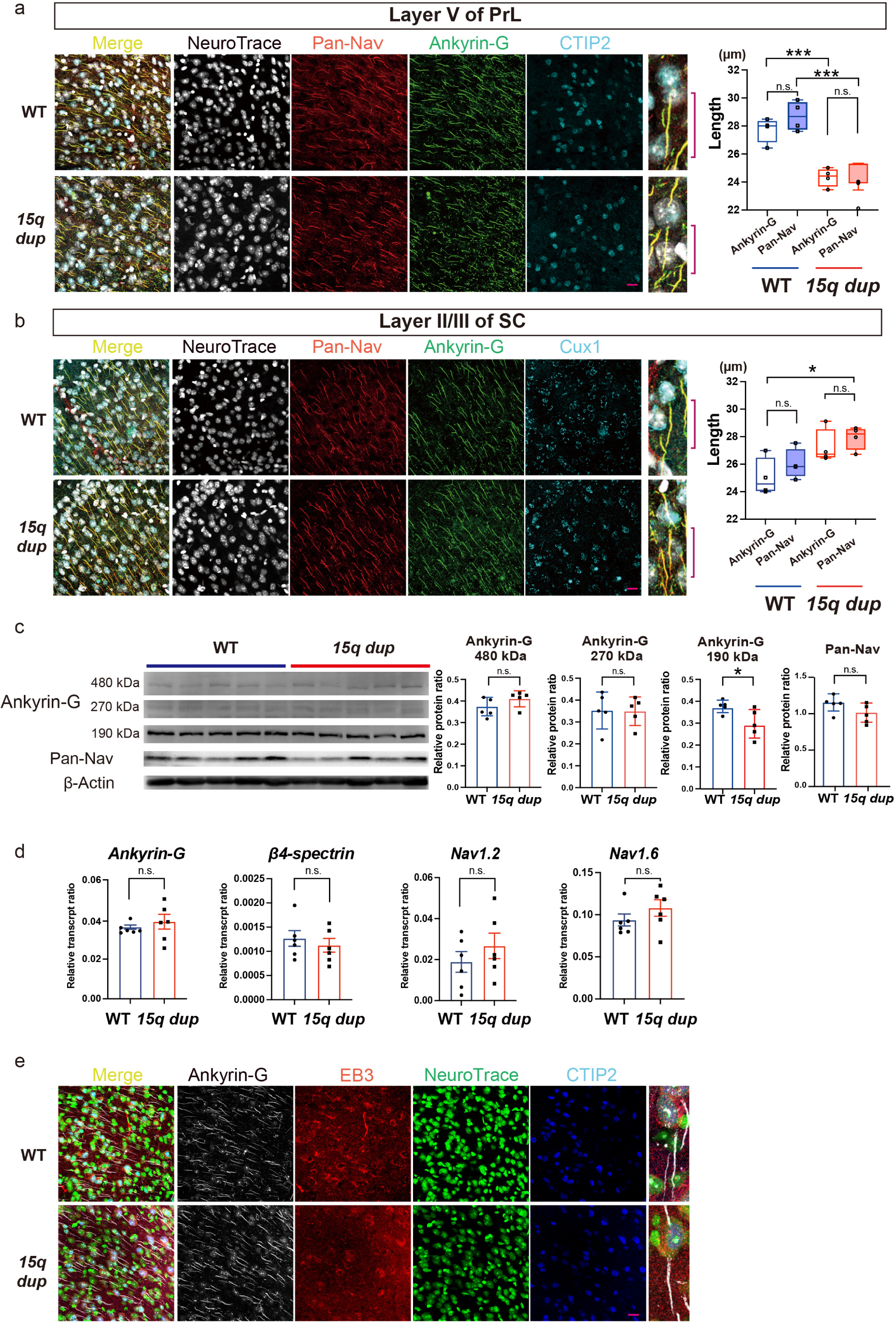
The alteration of axon initial segment (AIS) length of pyramidal neurons (PyNs) in *15q dup* mouse is not attributable to biochemical abnormality of Ankyrin-G and Pan-voltage-gated sodium (NaV) channels, aberrant accumulation of NaV channels, or end-binding protein 3 (EB3) at the AIS. **a)** Representative confocal images of Pan-NaV channels(red), Ankyrin-G(green), CTIP2(cyan), and NeuroTrace®(white) in layer V PyNs of the prelimbic cortex (PrL). Quantification of the length of Ankyrinn-G and Pan-NaV channels in layer V PyNs of two groups, wild-type (WT) and *15q dup* mice. Red bars indicate AIS. Scale bar = 10 µm. Values are presented as medians with 25–75% intervals, and error bars show the minimum to maximum values. n.s., not significant, left *** P=0.0009, right ***P=0.0003, analyzed with one-way analysis of variance (ANOVA) followed by post-hoc Tukey’s test (n = 4 per group). Scale bar, 10 µm. **b)** Representative confocal images of Pan-Nav(red), Ankyrin-G(green), Cux1(cyan), and NeuroTrace®(white) in layer II/III of the somatosensory cortex (SC). Quantification of the length of Ankyrin-G and Pan-NaV in layer II/III PyNs of two groups. Red bars indicate AIS. Scale bar = 10 µm. Values are presented as medians with 25–75% intervals, and error bars show the minimum to maximum values. n.s., not significant, *P=0.0192, analyzed with one-way ANOVA followed by post-hoc Tukey’s test (n = 4 per group). Scale bar, 10 µm. **c)** Representative images of western blot analysis of Ankyrin-G 480 kDa, 270 kDa, 190 kDa, Pan-NaV and β-actin. Quantification of relative protein ratio of Ankyrin-G and Pan-NaV protein levels compared to β-actin in two groups. All values are presented as the mean ± standard error of the mean (SEM). n.s., not significant; *P=0.0374, analyzed using two-tailed unpaired Student’s *t-*test (n = 6 per group). **d)** Quantification of relative transcript ratio of *Ankyrin G*, *β4-spectrin, Nav1.2*, and *Nav1.6* levels compared to *β-actin* in the whole cortex of two groups. All values are presented as the mean ± SEM. n.s., not significant, analyzed using two-tailed unpaired Student’s *t-*test (n = 6 per group). **E)** Representative confocal images of EB3(red), Ankyrin-G(white), CTIP2(blue), and NeuroTrace®(green) in layer V PyNs of the prelimbic cortex (PrL) in two groups. Scale bar = 10 µm.

In the same context, to demonstrate that AIS change in *15q dup* mice is based on circuit activity rather than biochemical changes in Ankyrin-G or Pan-NaV channels in the neuron, we quantified the protein levels of Ankyrin-G and Pan-NaV channels by western blot analysis in the PrL of WT and *15q dup* mice (Fig. 2c). We found that the levels of expression of 270 and 480 kDa Ankyrin-G, main AIS-related Ankyrin-G isoforms, and Pan-NaV channels did not significantly change in the PrL of *15q dup* mice (Ankyrin-G [480 kDa] protein ratio WT: 0.3741 ± 0.01913, *15q dup*: 0.4102 ± 0.01655; Ankyrin-G [270 kDa] protein ratio WT: 0.3524 ± 0.03773, *15q dup*: 0.3491 ± 0.02912; Pan-NaV channels protein ratio WT: 1.155 ± 0.05281, *15q dup*: 1.015 ± 0.05873). We also found decrease of 190 kDa Ankyrin-G, a dendrite-localizing Ankyrin-G isoform (Ankyrin-G [190 kDa] protein ratio WT: 0.3640 ± 0.01273, *15q dup*: 0.2857 ± 0.02874). Further, we examined transcript levels of AIS components, including *Ankyrin-G, β4-spectrin* and *NaV channels* using real-time polymerase-chain reaction (PCR) (*Ankyrin-G* transcript ratio WT: 0.03616 ± 0.001091, *15q dup*: 0.03890 ± 0.003732; *β4-spectrin* transcript ratio WT: 0.001266 ± 0.0001615, *15q dup*: 0.001125 ± 0.0001416; *NaV1.2* transcript ratio WT: 0.01889 ± 0.005055, *15q dup*: 0.02668 ± 0.006220; *NaV1.6* transcript ratio WT: 0.09373 ± 0.007036, *15q dup*: 0.1080±0.009804) (Fig. 2d). These results indicate that the changes in AIS length in *15q dup* mice are not attributable to biochemical changes in Ankyrin-G and NaV channels at the protein and transcript levels.

Furthermore, by examining EB3 accumulation in the AIS of both *15q dup* and WT mice, we investigated whether changes in the AIS length could be attributed to structural abnormalities. We observed no significant EB3 accumulation in the AIS of two groups (Fig. 2e), indicating that modifications to the AIS are likely dependent on neuronal activity.

### Alteration of electrophysiological properties in layer V pyramidal neurons in the prelimbic cortex of *15q dup* mice

To determine whether decreased AIS length corresponded to decreased neuronal excitability, we performed electrophysiological analysis with whole-cell patch-clamp recordings, using acute brain slices of the mPFC in WT and *15q dup* mice. We injected depolarizing current steps and found that the AP frequency was significantly decreased in *15q dup* mice (Fig. 3a). Consistent with these results, fʹ(Max), which is the maximum slope of the I-f curve was significantly decreased in *15q dup* mice (Fig. 3g). In addition, the frequency of spontaneous excitatory postsynaptic current (EPSCs) in layer V PyNs was significantly increased in the PrL of *15q dup* mice (Fig. 3e); however, there was no increase in current threshold, voltage threshold (Fig. 3c), resting membrane potential (RMP) (Fig. 3g), or the amplitude of EPSCs (Fig. 3e). All the electrophysiological data are shown in Table 1. These alterations were not observed in the IL (Figs. 3b, d, f, h). These results suggest that layer V PyNs in the PrL of *15q dup* ASD model mouse receive stronger response at the excitatory synapse and have reduced AP generation capacity.

**Fig. 3.**
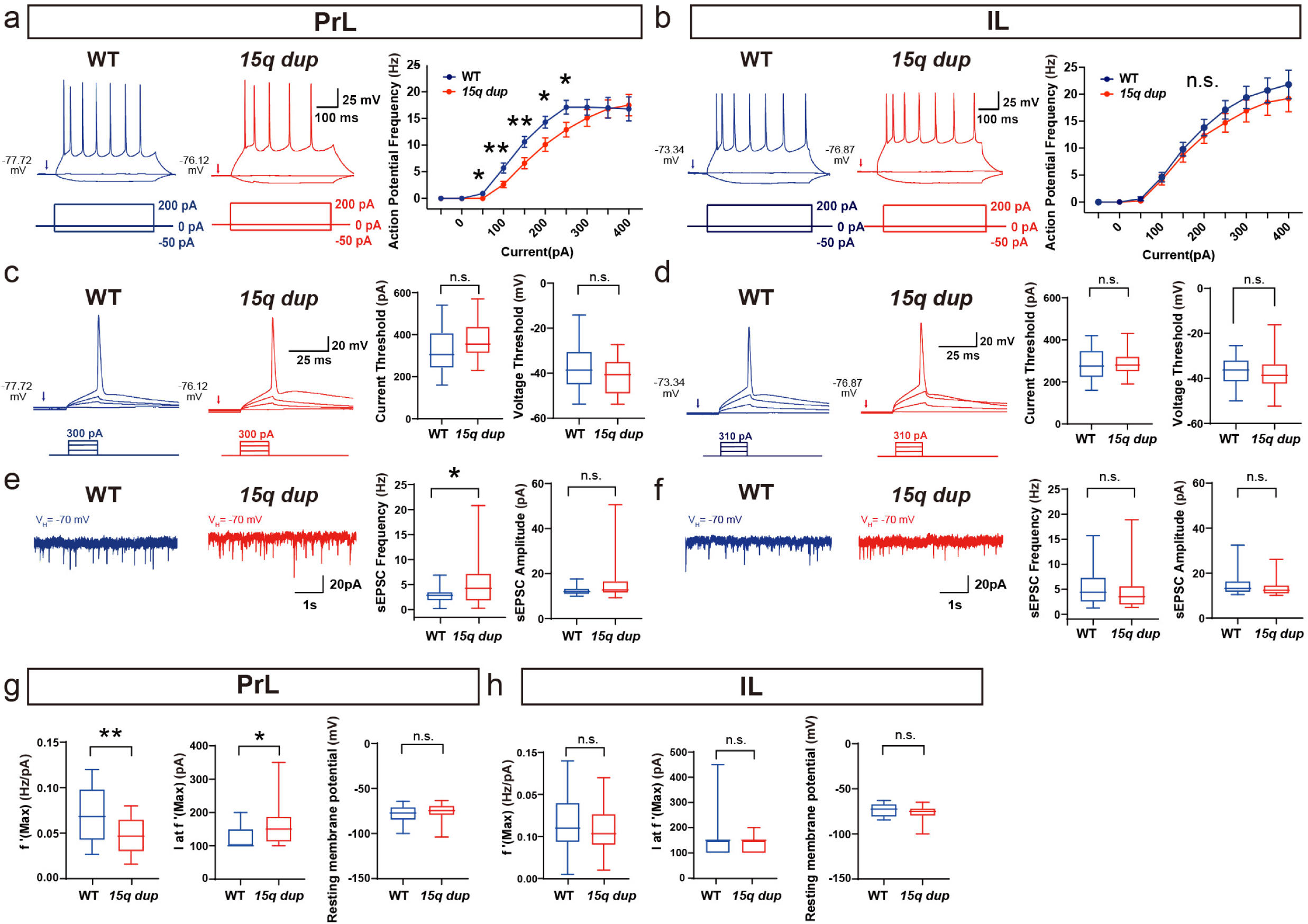
Specific alteration of electrophysiological properties in layer V pyramidal neurons (PyNs) in the prelimbic cortex (PrL) of *15q dup* mice. a, c, e, and g (left panels) show the analysis in PrL PyNs, and b, d, f, and h (right panels) show the analysis in the infralimbic cortex (IL). **a, b)** Representative traces of action potential (AP) trains evoked by current injection (500 ms, -50 pA and +200 pA). AP frequency was determined with 500 ms injections of indicated increasing currents. All values are presented as mean ± standard error of the mean (SEM). n.s., not significant, P values are from left * P=0.0396 (50 pA, PrL), ** P=0.0086 (100 pA, PrL), ** P=0.0093 (150 pA, PrL), * P=0.0167 (200 pA, PrL), * P=0.0167 (250 pA, PrL), analyzed with two-tailed unpaired Student’s *t-*test (n = 20 cells from at least 4 biologically independent experiments). **c, d)** Representative traces of single APs. The current and voltage threshold were measured with a step protocol of 20 ms pulses in 10 pA increments. All values are presented as median with 25–75% interval, and error bars show minimum to maximum values. n.s., not significant analyzed with two-tailed unpaired Student’s *t-*test (n = 20 cells from at least 4 biologically independent experiments). **e, f)** Representative traces of spontaneous excitatory postsynaptic currents (sEPSCs). The sEPSCs were recorded with the holding membrane at -70 mV in the voltage-clamp mode for 3 min. All values are presented as median with 25–75% interval, and error bars show minimum to maximum values. n.s., not significant, *P=0.0307, analyzed with two-tailed unpaired Student’s *t-*test (n = 20 cells from at least 4 biologically independent experiments). **g, h)** Analysis of maximum slope fʹ(max) of the I-f curve, the current at the maximum slope I at fʹ(max), and resting membrane potentials. All values are presented as median with 25–75% interval, and error bars show minimum to maximum values. n.s., not significant, * P=0.0345, ** P=0.0027, analyzed with two-tailed unpaired Student’s *t-*test (n = 20 cells from at least 4 biologically independent experiments).

**Table 1.**
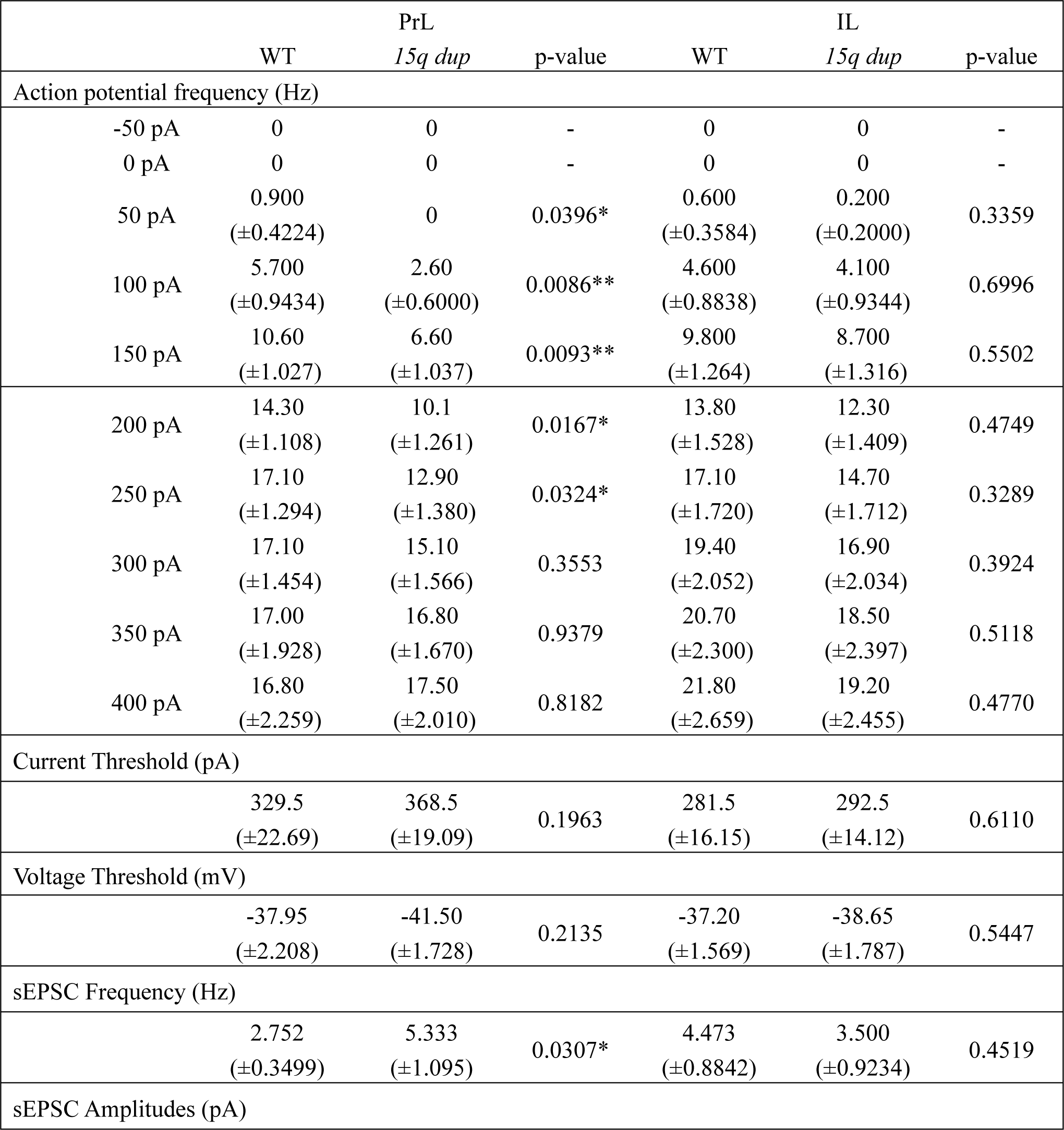
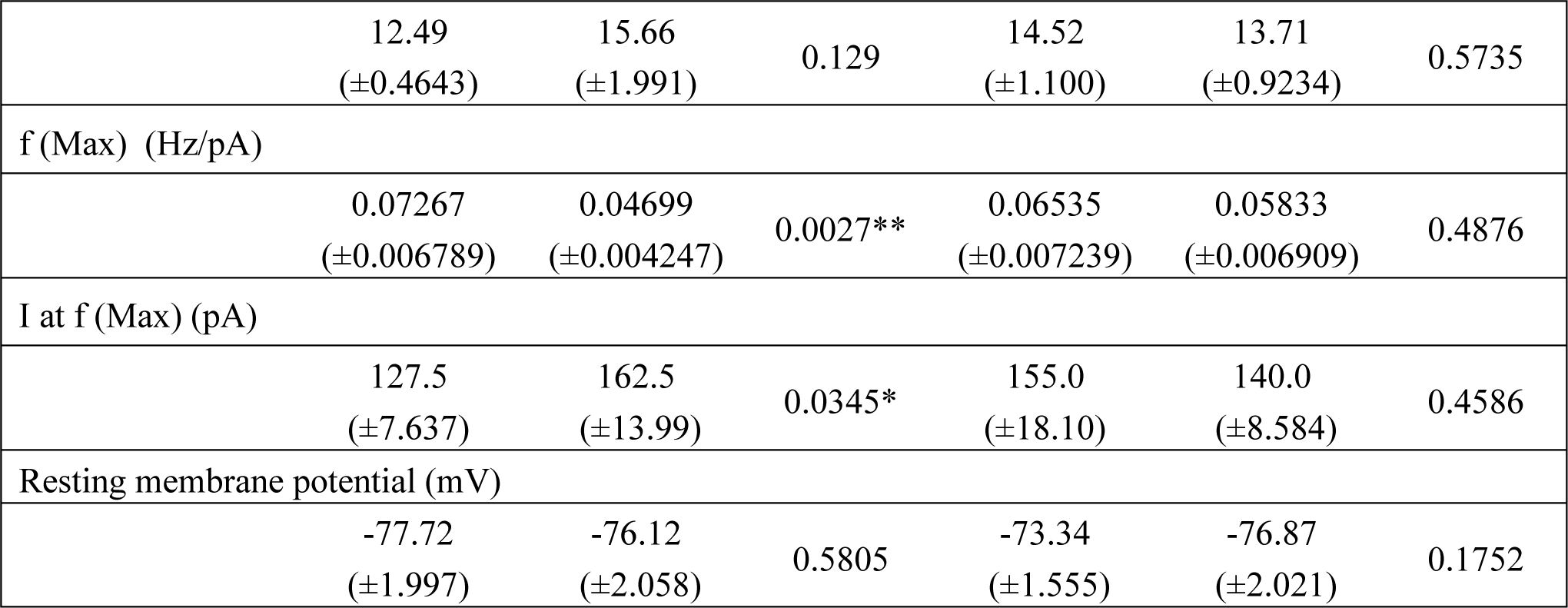
Electrophysiological data. The eight electrophysiological parameters are included in this examination and raw data were presented. The results of the analysis are shown in the images of **Fig. 3 a-h.** Prelimbic cortex is represented as PrL, Infralimbic cortex as IL, Wild Type as WT and 15q11-13 duplicaiton as *15q dup*. *p < 0.05, **p < 0.01, analyzed with student’s t test.

|  |  | PrL |  |  | IL |  |
| --- | --- | --- | --- | --- | --- | --- |
|  | WT | *15q dup* | p-value | WT | *15q dup* | p-value |
| Action potential frequency (Hz) |  |  |  |  |  |  |
| -50 pA | 0 | 0 | - | 0 | 0 | - |
| 0 pA | 0 | 0 | - | 0 | 0 | - |
| 50 pA | 0.900<br>(±0.4224) | 0 | 0.0396* | 0.600<br>(±0.3584) | 0.200<br>(±0.2000) | 0.3359 |
| 100 pA | 5.700<br>(±0.9434) | 2.60<br>(±0.6000) | 0.0086** | 4.600<br>(±0.8838) | 4.100<br>(±0.9344) | 0.6996 |
| 150 pA | 10.60<br>(±1.027) | 6.60<br>(±1.037) | 0.0093** | 9.800<br>(±1.264) | 8.700<br>(±1.316) | 0.5502 |
| 200 pA | 14.30<br>(±1.108) | 10.1<br>(±1.261) | 0.0167* | 13.80<br>(±1.528) | 12.30<br>(±1.409) | 0.4749 |
| 250 pA | 17.10<br>(±1.294) | 12.90<br>(±1.380) | 0.0324* | 17.10<br>(±1.720) | 14.70<br>(±1.712) | 0.3289 |
| 300 pA | 17.10<br>(±1.454) | 15.10<br>(±1.566) | 0.3553 | 19.40<br>(±2.052) | 16.90<br>(±2.034) | 0.3924 |
| 350 pA | 17.00<br>(±1.928) | 16.80<br>(±1.670) | 0.9379 | 20.70<br>(±2.300) | 18.50<br>(±2.397) | 0.5118 |
| 400 pA | 16.80<br>(±2.259) | 17.50<br>(±2.010) | 0.8182 | 21.80<br>(±2.659) | 19.20<br>(±2.455) | 0.4770 |
| Current Threshold (pA) |  |  |  |  |  |  |
|  | 329.5<br>(±22.69) | 368.5<br>(±19.09) | 0.1963 | 281.5<br>(±16.15) | 292.5<br>(±14.12) | 0.6110 |
| Voltage Threshold (mV) |  |  |  |  |  |  |
|  | -37.95<br>(±2.208) | -41.50<br>(±1.728) | 0.2135 | -37.20<br>(±1.569) | -38.65<br>(±1.787) | 0.5447 |
| sEPSC Frequency (Hz) |  |  |  |  |  |  |
|  | 2.752<br>(±0.3499) | 5.333<br>(±1.095) | 0.0307* | 4.473<br>(±0.8842) | 3.500<br>(±0.9234) | 0.4519 |
| sEPSC Amplitudes (pA) |  |  |  |  |  |  |
|  | 12.49<br>(±0.4643) | 15.66<br>(±1.991) | 0.129 | 14.52<br>(±1.100) | 13.71<br>(±0.9234) | 0.5735 |
| f (Max) (Hz/pA) |  |  |  |  |  |  |
|  | 0.07267<br>(±0.006789) | 0.04699<br>(±0.004247) | 0.0027** | 0.06535<br>(±0.007239) | 0.05833<br>(±0.006909) | 0.4876 |
| I at f (Max) (pA) |  |  |  |  |  |  |
|  | 127.5<br>(±7.637) | 162.5<br>(±13.99) | 0.0345* | 155.0<br>(±18.10) | 140.0<br>(±8.584) | 0.4586 |
| Resting membrane potential (mV) |  |  |  |  |  |  |
|  | -77.72<br>(±1.997) | -76.12<br>(±2.058) | 0.5805 | -73.34<br>(±1.555) | -76.87<br>(±2.021) | 0.1752 |

### Alteration of AIS plasticity in layer V PyNs in the mPFC of *15q dup* mice by depolarization with potassium chloride

We examined AIS plasticity in acute slices of the mPFC by depolarization with high-concentration potassium chloride (KCl) *in vitro*. As shown in Fig. 4, we found that a 1-hour bath with high-concentration KCl was sufficient to shorten the AIS length of layer V PyNs in the PrL and IL of WT mice, and the normal AIS length was restored within 3 hours. To determine whether this phenomenon was maintained in *15q dup* mice, we quantified the AIS length in mPFC acute slices from *15q dup* mice. The AIS length in the PrL of *15q dup* mice was decreased even before KCl application (0 hour), and high-concentration KCl exposure did not change the AIS length after 1 and 3 hours of exposure (WT 0 hr: 29.18 ± 1.409 μm; 1 hr: 25.44 ± 0.6167 μm; 3 hr: 29.24 ± 1.088 μm; *15q dup* 0 hr: 26.72 ±0.2752 μm; 1 hr 25.22 ±0.8425 μm; 3 hr: 24.46 ±0.8334 μm) (Fig. 3b). In addition, the AIS length of IL PyNs of *15q dup* mice showed only impaired AIS plasticity (WT 0 hr: 27.20 ±0.8161 μm; 1 hr: 23.08 ±0.7186 μm; 3 hr: 26.74 ±0.9855 μm; *15q dup* 0 hr: 25.68 ±0.6064μm; 1 hr 24.01 ±0.6707 μm; 3 hr: 25.29 ±1.396 μm) (Fig. 3d). These results indicate that while abnormal AIS plasticity of layer V PyNs broadly occurred in the PrL and IL of *15q dup* ASD model mice, AIS shortening occurred specifically in the layer V PyNs in the PrL.

**Fig. 4.**
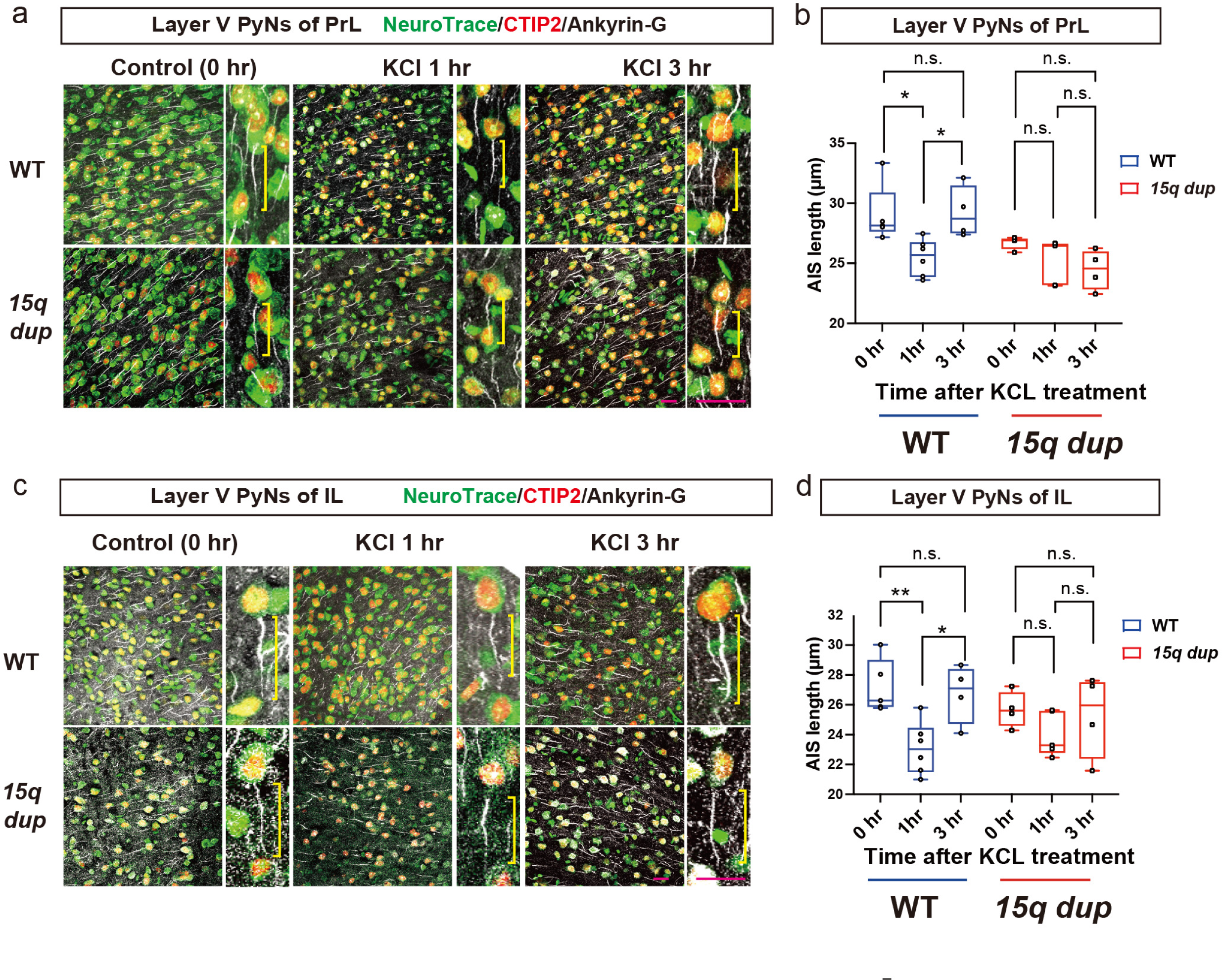
Alteration of axon initial segment (AIS) plasticity in layer V pyramidal neurons (PyNs) in the medial prefrontal cortex (mPFC) of *15q dup* mice. a and b (upper panels) show the analysis in the prelimbic cortex (PrL) PyNs, while c and d (lower panels) show the analysis in the infralimbic cortex (IL). **a, c)** Representative confocal images of the layer V PyNs in the PrL (**a**) or IL (**c**) of wild-type (WT) and *15q dup* mice stained with NeuroTrace®(green), anti-CTIP2(red) and anti-Ankyrin-G(white) antibodies. Brain slices were treated with high-concentration potassium chloride (8 mM) for indicated time. 0 hr (control), 1 hr, or 3 hr. Left * P=0.0110, right * P=0.0118, **b, d)** Quantification of the AIS length of the layer V PyNs in the PrL (**a**) or IL (**c**) of wild-type (WT) and *15q dup* mouse brain slices. All values are presented as median with 25–75% interval, and error bars show minimum to maximum values with all data points. n.s., not significant, * P=0.0185, ** P=0.0044, analyzed with two-way analysis of variance followed by post-hoc Sidak’s multiple comparison test (n = 4–6 per group). Scale bar, 10 µm. Yellow bars indicate the AIS length of each neuron.

### AIS length of PyNs projecting to specific areas are significantly decreased in *15q dup* mice

To assess the AIS shortening is dependent on the local excitatory connections^35^, or neural circuit targets, we aimed to determine whether long-distance circuit-specific AIS abnormalities can be detected using the retrograde tracing method. Previous reports have shown that layer V PyNs of the PrL project to various brain areas, including the nucleus accumbens (NAcc), ventromedial caudate-putamen (CPVM), basolateral amygdala (BLA), lateral habenula (LHb), ventral tegmental area (VTA), and dorsal raphe nucleus (DRN)^36^. In our study, we stereotaxically injected the cholera toxin B subunit (CTB) into these targeted areas and confirmed the presence of projections in the PrL (Fig. 5a, b). Our results indicated that layer II/III superficial PyNs predominantly project to the BLA (Fig. 5 a, b); thus, we quantified five target areas (NAcc, CPVM, LHb, VTA, and DRN). In addition to the diversity of projecting neurons with respect to AIS length, there was significant decrease in the AIS length of PyNs projecting to the NAcc (WT: 28.72 ±0.4104 μm; *15q dup*: 26.46 ±0.6579 μm), LHb (WT: 27.50 ±0.4405 μm; *15q dup*: 24.08 ±0.7046 μm), and DRN (WT: 28.20 ±0.5685 μm; *15q dup*: 25.62 ±0.3164 μm), but not of those projecting to the CPVM (WT: 26.95 ±0.9346 μm; *15q dup*: 26.87 ±0.8498 μm), and VTA (WT: 26.37 ±1.206 μm; *15q dup*: 26.00 ±1.370 μm) (Fig 5c). As shown in Fig. 5d, the cumulative frequency plots of individual AIS lengths in WT and *15q* mice showed a leftward shift in the PyNs projecting to the NAcc, LHb, and DRN, no rightward shift, and no significant shift in the neurons projecting to the CPVM and VTA, as observed in Fig. 5c. Integrating all these results, we conclude that abnormal AIS length can signal target-specific neural circuit abnormalities in *15q dup* mice.

**Fig. 5.**
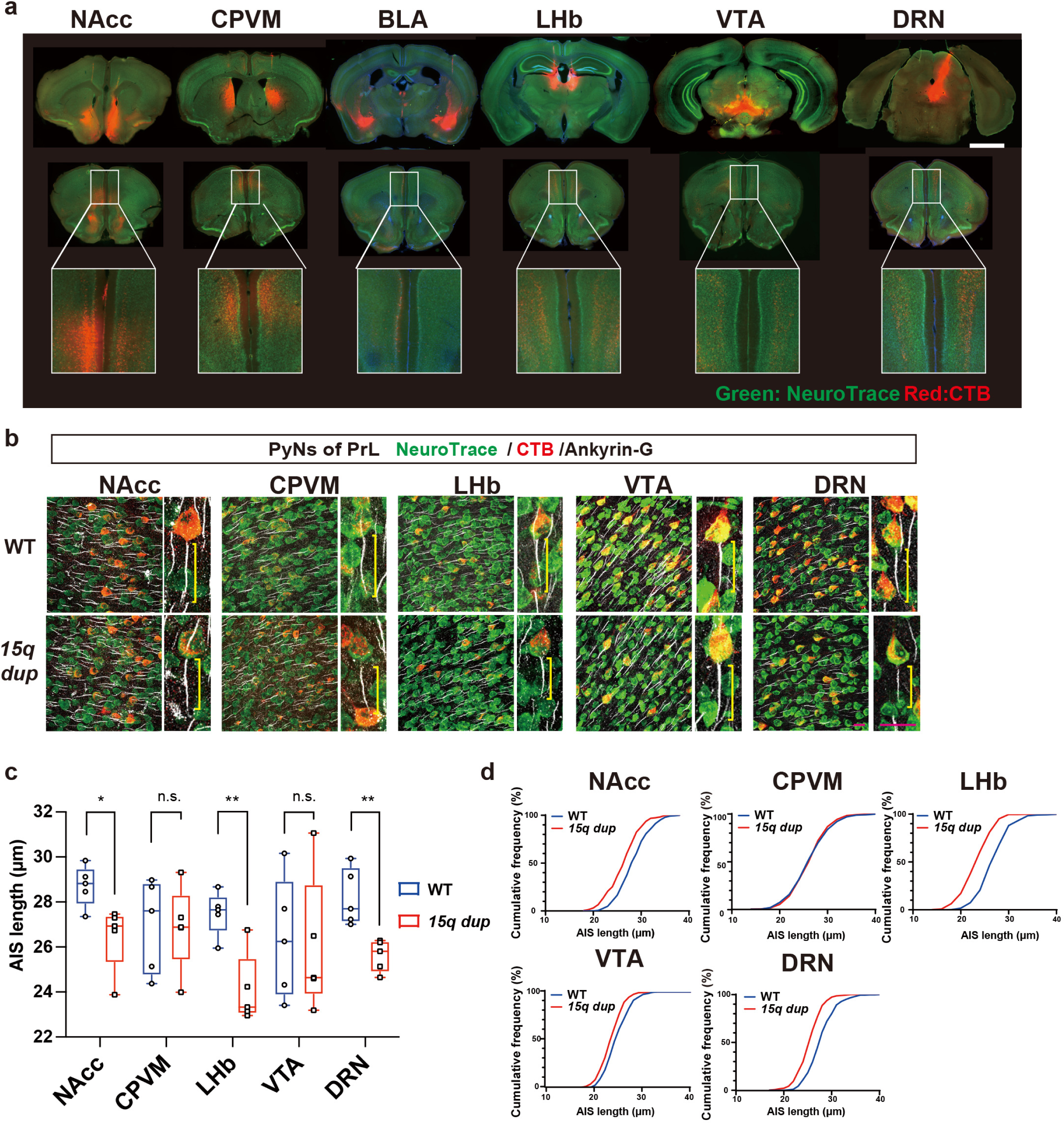
The length of axon initial segment (AIS) of layer V pyramidal neurons (PyNs) in the prelimbic cortex (PrL) of *15q dup* mice varies depending on the projection target. **a)** Upper: Representative retrograde tracing images of NeuroTrace® (green) and cholera toxin B subunit (CTB) (red). CTB was injected into the nucleus accumbens (NAcc), caudate putamen, ventromedial division (CPVM), lateral habenula (LHb), ventral tegmental area (VTA), and dorsal raphe nucleus (DRN). Lower: Representative image of the projected area corresponding to the medial prefrontal cortex (mPFC). Scale bar= 100 µm. **b)** Representative confocal images of the axon initial segment (AIS) and cumulative frequency of AIS length in layer V PyNs of wild-type (WT) and *15q11-13* duplication (*15q dup*) mice. Fluorescent colors indicate NeuroTrace®(green), CTB(red), and Ankyrin-G(white). Scale bar= 10 µm. **c)** Analysis of AIS in layer V PyNs in the mPFC projecting to the NAcc, CPVM, LHb, VTA, and DRN in WT and *15q dup* mouse brain. All values are presented as median with 25–75% interval, and error bars show minimum to maximum values with all data points, * P=0.0195 (NAcc), left ** P=0.0034 (LHb), right ** P=0.0041 (DRN), analyzed with two-tailed unpaired Student’s *t-*test (n = 5 per group). **d)** Analysis of the cumulative frequency of AIS length in layer V PyNs in the mPFC projecting to the NAcc, CPVM, LHb, VTA, and DRN in WT(blue) and *15q dup*(red) mouse brains.

## DISCUSSION

In our investigation, we aimed to identify functionally aberrant AIS structures that can serve as biomarkers of task-independent neural activity in ASD model mice. We examined the cortical regions and layers for abnormalities in the AIS length of PyNs in *15q dup* ASD model mice. We observed a decrease in AIS length in layer V PyNs of the PrL and an elongation in the neurons of the SC in these model mice, without any task. These results suggest region-specific alterations in AIS, with potential implications for neural circuit function and ASD pathophysiology. Our comprehensive analysis excluded biochemical perturbations in Ankyrin-G, misplaced NaV channels, and aberrant accumulation of EB3 as causes of these AIS changes. Therefore, these modifications may reflect aberrant task-independent neural activity. Electrophysiological studies using whole-cell patch-clamp recordings in the PrL revealed an increase in the frequency of spontaneous excitatory postsynaptic currents (sEPSCs) and a diminished propensity for APs generation upon current injections into the PrL. Subsequently, we discovered deficient AIS plasticity in the PrL and IL when subjected to high-concentration KCl conditions in the *15q dup* mice. Additionally, retrograde tracing revealed that AIS shortening was dependent on the target projection.

The first question that arises is, what triggers the specific alteration in the length within the mPFC and SC of *15q dup* mice without any task? Prior investigations have indicated a deficiency in inhibitory synapses originating from serotonergic neurons in the primary somatosensory cortex (S1BF), leading to over-excitation of layer II/III PyNs^27^. This hyperexcitability in the SC may mirror the sensory-related manifestations observed in individuals with ASD^22^. Our findings support this hypothesis by demonstrating that hyperactive SC neurons have extended AIS.

The widespread impact of serotonergic fibers prompts the question of why PyNs in the PrL exhibit specific inactivation in contrast to the activation seen in SC PyNs. Despite the elevated levels of sEPSCs, PrL neurons exhibit reduced AP generation in response to current injections. This paradoxical phenomenon may be explained by “homeostatic adaptation,” a mechanism by which neurons adjust their excitability to maintain network stability^37^. Currently, there are three main compartments for this adaptation: at the AIS, excitatory and inhibitory postsynaptic compartments, and presynaptic boutons^37^. The exact mechanism by which PrL neurons adapt to sustainably reduced inhibitory serotonergic inputs remains unclear and requires further investigation, especially regarding changes in the AIS.

Conversely, *Tau* mutations linked to frontotemporal dementia have been found to restrict AIS plasticity through an EB3-dependent mechanism, leading to a pathologically static AIS^21^. Neurons derived from patient-specific induced pluripotent stem cells with *Tau* mutations showed reduced AIS length without affectation of AIS plasticity under high-concentration-potassium treatment. Our study also confirmed that AIS reduction in *15q dup* mice was not due to abnormal EB3 accumulation, as evidenced by comparable EB3 and Ankyrin-G immunostaining between WT and *15q dup* mice. Regarding structural AIS alterations, slight reduction in the 190 kDa Ankyrin-G isoform localized to dendrites was observed in the brains of *15q dup* mice brain, while the principal AIS-associated isoforms (270 kDa and 480 kDa) remained unaltered^38^, indicating that abnormal synaptic transmission at dendrites may be related to post-translational protein modifications, such as palmitoylation^39^. Intriguingly, our results delineated two distinct neuronal states: a pathologically static AIS state that is dependent on EB3-mediated structural changes^21^ and an activity-dependent static AIS state that is not linked to abnormal EB3. Future applications of designer receptors exclusively activated by designer drugs (DREADD) or optogenetics will clarify whether these AIS alterations are a consequence of activity-dependent mechanisms or represent irreversible structural changes.

In the context of the *15q dup* ASD model, we detected abnormal AIS in the PyNs projecting from the PrL to the NAcc, LHb, and DRN, but not to the VTA, CP, or BLA. To discern the neural circuits associated with aberrant behaviors in this ASD model, techniques such as DREADD or optogenetics have been employed to selectively activate neurons in layer V. Considering the variation in neural circuits involved in social behavior across ASD models^28^, a comprehensive understanding of these neural networks is essential for identifying atypical neural circuits through potential circuit-based biomarkers in each model. For neural circuit mapping during specific tasks, researchers have developed methods to capture neural activity snapshots via immediate early genes such as *Fos* or *Arc*, monitoring intracellular Ca^2+^ concentration, and recording synaptic events^32, 33^.

This study raises a critical question: How does AIS analysis diverge from the other techniques discussed? AIS analysis may provide a unique perspective on spontaneous intrinsic neural network functions, such as those of the default mode network (DMN), which exhibit increased activity during various cognitive processes, including those related to social cognition^40^. In contrast, the DMN exhibits decreased activity during tasks that require focused attention^41^. The vulnerability of the DMN to disruptions in ASD^42^ and other neuropsychiatric conditions such as Alzheimer’s disease, depression, and schizophrenia^40^ has recently been highlighted, with the network playing a consistent role in the processing of social information and contexts. It encompasses the understanding of mental states, deducing others’ thoughts, and forming self-referential narratives^43^. Recent advances in imaging and tracing methodologies have elucidated the DMN architecture in mice, identifying PrL and IL as central components through resting-state functional magnetic resonance imaging and neural tracing^44^. Previous investigations in *15q dup* mice also corroborated these findings, revealing abnormalities in the DMN^30^. Further human studies in this regard are essential; however, our results imply that AIS abnormalities may reflect underlying neuronal pathologies and could serve as biomarkers for defective spontaneous neural circuits, particularly in the task-independent neural activities of individuals with neuropsychiatric disorders.

In summary, AIS analysis provides insights into the structural integrity of neurons, thereby providing a comprehensive perspective on the neural network dysfunction that is characteristic of neuropsychiatric conditions, including ASD.

## METHODS

### Materials

The chemicals used in this study were of the highest purity and obtained from Fujifilm Wako Pure Chemical Co. (Osaka, Japan), Sigma–Aldrich Japan/Merck (Tokyo, Japan), and Nacalai Tesque (Kyoto, Japan).

### Mice

*15q* dup mice were generated using a chromosome engineering technique^26^. They were maintained on a C57BL/6J background. Genotyping PCR of mouse tail genomic DNA was performed as described previously^26^, with some modifications. Briefly, genotyping PCR was performed using Go Taq Green Master Mix (Promega, Madison, WI, USA) with the following four primer sets: hprt-intron2-F1: AGAGGAGGGCCTTACTAATTACTTA; hprt-intron2-R2: ATATGTACTTTTGCATATAGTATAC; olMR0015: AAATGTTGCTTGTCTGGTG; olMR0016: GTCAGTCGAGTGCACAGTTT. PCR conditions were as follows: pre-denaturation at 95°C for 2 min, followed by 30 amplification cycles of denaturation at 95°C for 30 sec, primer annealing at 58°C for 30 sec, and extension at 72°C for 30 sec, with a final additional extension at 72°C for 5 min. Littermates or age-matched WT mice were used as controls. Mouse lines were maintained in a transgenic mouse room in a specific pathogen-free environment at the Department of Experimental Animals, Interdisciplinary Center for Science Research, Head Office for Research and Academic Information, Shimane University, under the university guidelines for the care and use of animals. All the animal protocols were approved by the Department of Experimental Animals, Interdisciplinary Center for Science Research, Head Office for Research and Academic Information, Shimane University (approval number: IZ4-4).

### Immunofluorescence study

Immunostaining was performed as described previously^45^. Summarily, brains were obtained from anesthetized 8-week-old mice that were injected intraperitoneally with an anesthetic combination that comprised 0.3 mg/kg medetomidine, 4.0 mg/kg midazolam, and 5.0 mg/kg butorphanol, and then subjected to cardiac perfusion fixation using 4% paraformaldehyde (PFA) in 0.01 M phosphate buffer saline (PBS; pH 7.4) for Ankyrin-G and 3% glyoxal/0.8 acetic acid in 0.1 M phosphate buffer for Pan-NaV channel. After cardiac perfusion fixation, post-fixation was performed for 2 h (PFA) or 24 h (glyoxal). Post-fixed brains were rinsed with saline and cryoprotected overnight with 30% sucrose in PBS at 4°C. Brain sections (50 μm) were cut using a sliding microtome (HM430; Thermo Fisher Scientific, Waltham, MA, USA ). Brain slices were pre-incubated with 0.01 M PBS (pH 7.4) containing 0.4% Triton X-100 and 3% normal donkey serum (PBTDS) for 1 h. Afterwards, the samples were incubated two overnight at 4°C with primary antibodies diluted to appropriate concentrations in PBTDS. Samples were thoroughly rinsed in PBST, followed by application of fluorescently labeled secondary antibodies overnight at 4°C. Finally, fluorescent-labeled samples were rinsed with 0.01 M PBST every 5 min. Samples were placed on gelatin-coated glass slides and cover glasses (Matsunami Glass, Osaka, Japan) using Vectashield (Vector Laboratories, Burlingame, CA, USA), and examined using FV-1000D and FV-3000D confocal microscopes (Olympus, Tokyo, Japan). The antibodies and reagents used are listed in Table 2.

**Table 2.**
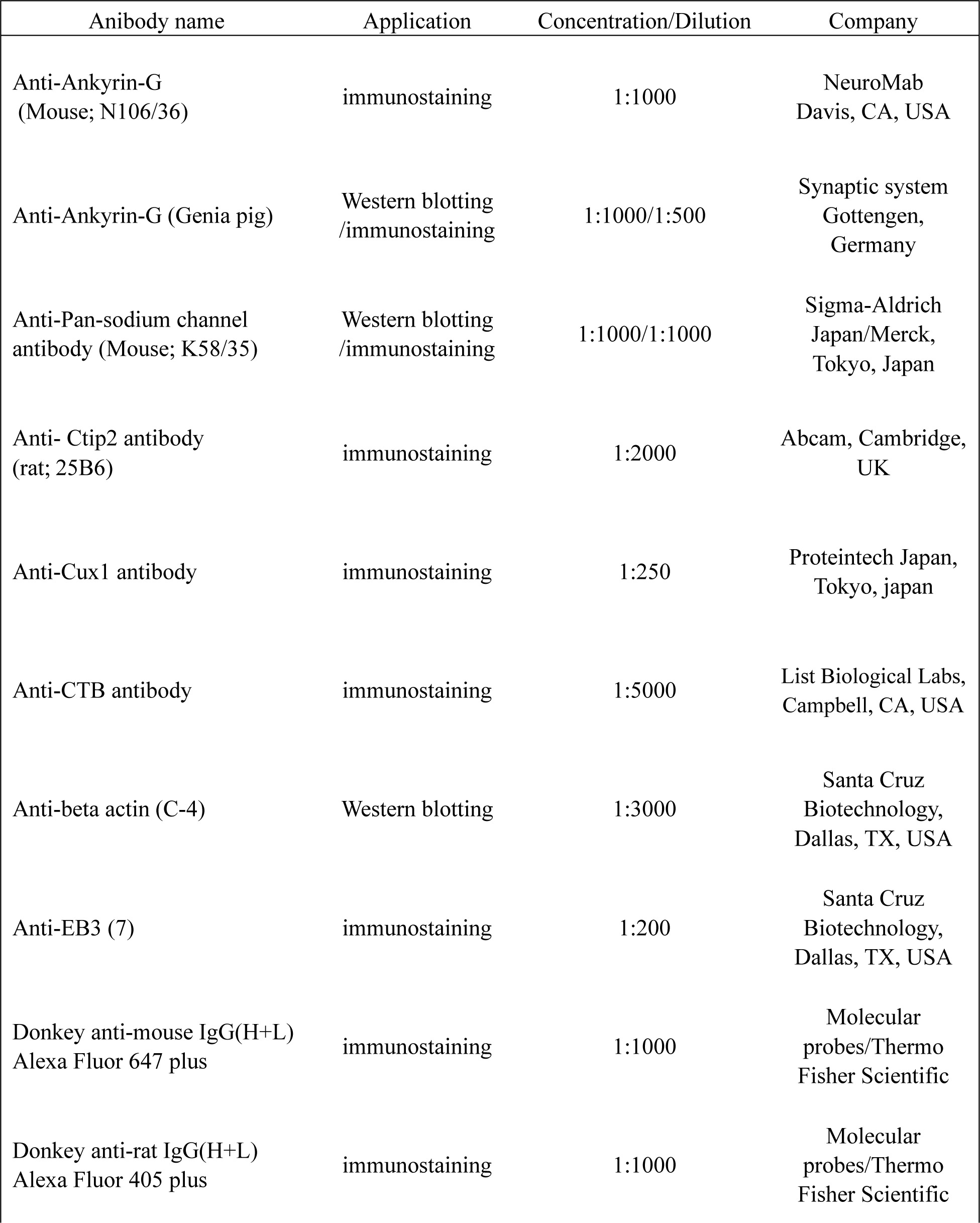
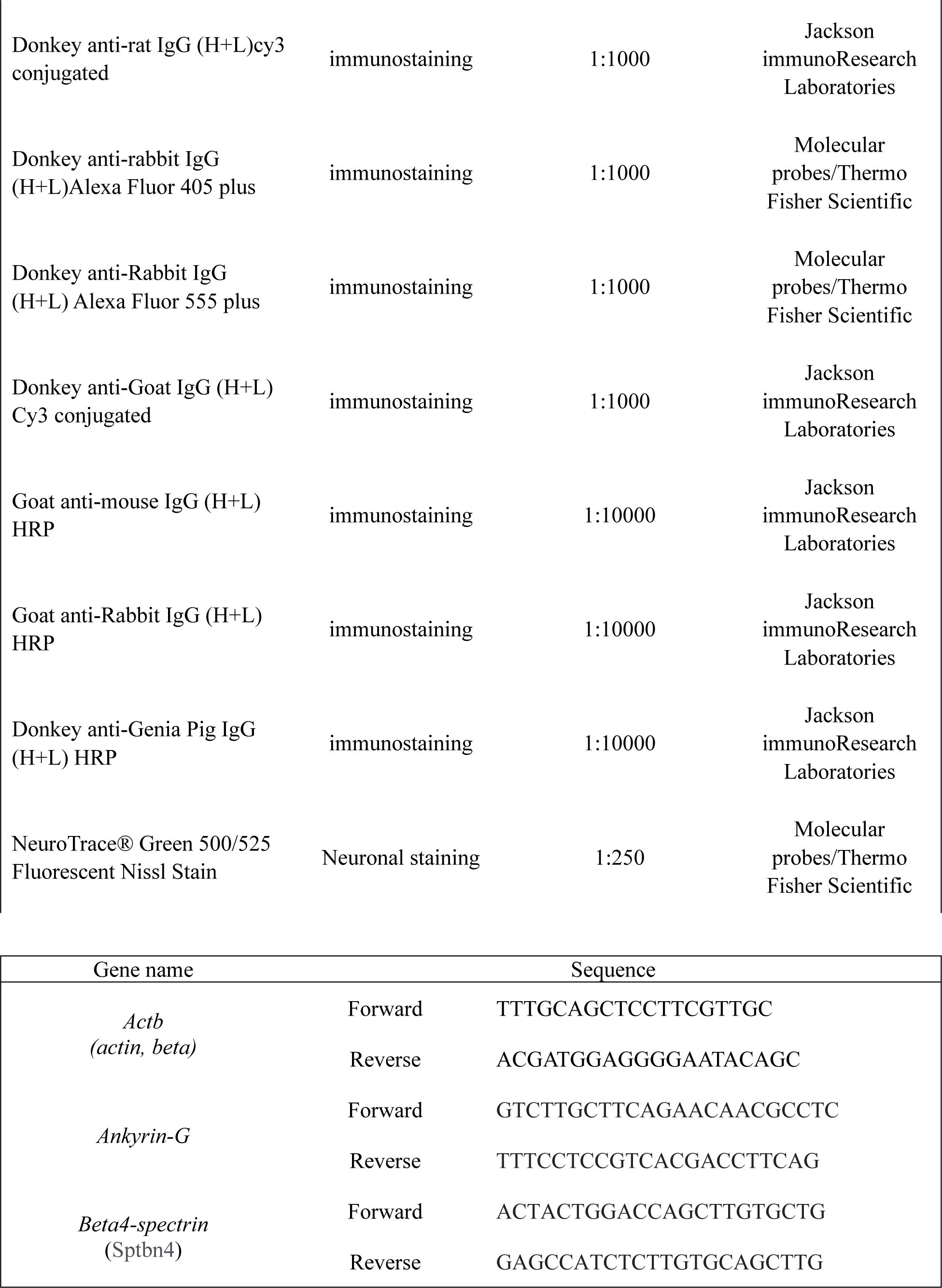
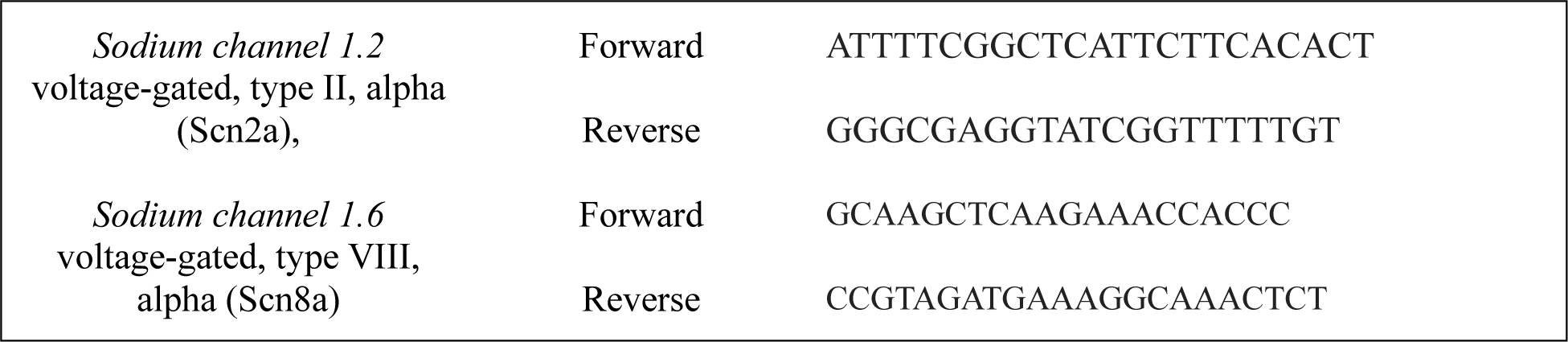
Antibody, qPCR primer and reagent list.

### Electrophysiology

Mice brains were quickly harvested and placed in ice-cold artificial cerebrospinal fluid (ACSF) containing 124 mM NaCl, 2.5 mM KCl, 2 mM CaCl_2_, 1 mM MgSO_4_, 25 mM NaHCO_3_, 1 mM NaH_2_PO_4_, and 10 mM glucose saturated with carbogen (95% O_2_, 5% CO_2_). Coronal brain slices (300 mm), including the mPFC region, were prepared using a vibratome (7000smz-2, Campden, Loughborough, Leics, England)^46^. Brain slices were placed in a chamber filled with carbogen-saturated ACSF at room temperature for approximately 1 h. All experiments were performed in a recording chamber on the stage of a BX51WI microscope (Olympus, Center Valley, PA, USA), using infrared differential interference contrast optics for neuronal visualization. Layer V pyramidal neurons in the PrL or IL were visually identified and whole-cell patch-clamp recordings were performed using an amplifier (Axopatch 200 B, Molecular Devices, San Jose, CA, USA) at room temperature. Recording pipettes were pulled from borosilicate glass (outer diameter 1.5 mm, inner diameter 1.12 mm, World Precision Instruments, Sarasota, FL, USA) to a tip resistance of 2-5 MΩ. In the current-clamp or voltage-clamp modes, tip microelectrodes were filled with internal liquid comprising of 120 mM K-gluconate, 5 mM NaCl, 1 mM MgCl_2_, 0.5 mM ethylene glycol tetra-acetic acid (EGTA), 2 mM Mg-ATP, 0.1 mM Na_3_GTP, and 10 mM HEPES; pH 7.2, 280-300 milliosmole were used for measuring RMPs, APs, and sEPSCs. The RMP was measured directly upon entering the whole-cell configuration in the current clamp at I = 0. The AP properties were measured with a step protocol of 20 ms pulses in 10 pA increments, starting from a holding current of I = 0. For firing pattern analysis (including the I-f curves), 500 ms pulses incrementing in 50 pA steps were used to trigger AP trains. The maximum slope of the curve and current at the maximum slope were calculated for each group. To record sEPSCs, the holding membrane was maintained at -70 mV in voltage-clamp mode for 3 min. APs and sEPSCs were analyzed using Mini Analysis Software, and membrane potentials were analyzed using Clampfit 10.7. All electrophysiological parameters are shown in Table 1.

### AIS plasticity analysis

The acute brain sections were prepared using the same method as described above. Coronal brain slices (300 μm) were put in a chamber for storing slices filled with carbogen-saturated ACSF at 37°C with or without 8 mM KCl for depolarization for 1 and 3 hours. After depolarization, brain slices were post-fixed in 4% PFA for 30 min on ice. Post-fixed brains were rinsed with saline and then cryoprotected overnight with 30% sucrose in PBS at 4°C. Next, brain sections (50 μm) were prepared using a cryostat (CM1900) (Leica, Wetzlar, Germany), placed on FRONTIER glass slides (Matsunami Glass, Osaka, Japan), and stored at -20°C until use. Brain slices were pre-incubated with 0.01 M PBS (pH 7.4) containing PBTDS for 1 h. Afterwards, samples were incubated 40 hours at 4°C with primary antibodies diluted to appropriate concentrations in PBTGS. Samples were thoroughly rinsed in PBST, followed by overnight application of fluorescently labeled secondary antibodies at 4°C. Finally, fluorescent-labeled samples were rinsed with 0.01 M PBST every 5 min. Samples were mounted on cover glasses using Vectashield (Vector Laboratories) and analyzed using FV-1000D and FV-3000D confocal microscopy (Olympus).

### Western blotting

mPFC homogenates were prepared from *15q dup* and WT mice. All procedures were performed on ice or at 4°C. mPFC was homogenized using an Ultrasonic Homogenizer SONIFIER 450 Advanced (BRANSON, Danbury, USA) in radio immunoprecipitation assay buffer (WAKO, Osaka, Japan) containing 2 mM EGTA and Protease Inhibitor Cocktail Set III (Wako, Osaka, Japan). To remove chromosomal DNA, cell debris, and fibers, the homogenates were centrifuged at 800 × g for 10 min, and supernatants were collected and stored as whole homogenate fractions at -80°C. Protein concentrations were determined using bicinchoninic acid assay (Thermo Fisher Scientific). Samples were subjected to sodium dodecyl-sulfate polyacrylamide gel electrophoresis and immunoblot analyses using various antibodies.

Western blotting was performed as described previously^45^ with minor modifications. Briefly, each sample was separated on 5-20% gradient e-PAGEL mini gels (ATTO Corporation, Tokyo, Japan) and transferred to polyvinylidene fluoride membranes (Merck Millipore, Billerica, MA, USA). The membranes were incubated for 30 min with blocking buffer (Nacalai Tesque). They were then incubated overnight with primary antibodies diluted in 5% Blocking One in tween-tris-buffered saline (T-TBS) buffer and washed thrice with T-TBS prior to incubation for 1 h with secondary antibodies in T-TBS. After washing thrice in T-TBS, immunoreactivity was detected using the Enhanced chemiluminescence (ECL) system (GE Healthcare, Tokyo, Japan). Both chemiluminescence and pre-stained size marker images were captured in the same field of view using ImageQuant 800 (AMERSHAM/GE Healthcare Japan, Tokyo, Japan). For quantification, band intensities were measured using ImageJ software (https://imagej. net/welcome). Full images of western blots are shown in Supplementary Fig. 4. The antibodies used are listed in Table 2.

### RNA extraction and quantitative real-time PCR

RNA extraction and quantitative real-time PCR were performed as described previously with modifications^45^. Briefly, the mouse brains were dissected and snap-frozen in liquid nitrogen. The frozen tissue was ground into powder using a BioMasher II (Nippi, Tokyo, Japan). Total RNA was extracted using Isogen II (Nippon Gene, Tokyo, Japan). The mRNA of various genes were quantified using a fluorescence-based real-time detection method. First-strand cDNA was synthesized using ReverTra Ace qPCR RT Master Mix with gDNA Remover (Toyobo, Osaka, Japan). Real-time PCR was performed using a Thermal Cycler Dice Real-Time System II TP900 (Takara Bio, Shiga, Japan), and the relative copy number of various gene transcripts per beta-actin transcript was determined using calibration standards for each tested molecule. The primers used are listed in Table 2.

### Retrograde Tracing

Injections of CTB (List Biological Labs, Campbell, CA) into the NAcc (A: 1.30, L: ±0.85, D: -3.80), CPVM (A: 0.26, L: ±1.25, D: -3.0), LHb (A: -1.4, L: ±0.75, D: -3.90), VTA (A: -3.3, L: ±0.4, D: -4.0), and DRN (A: -2.24, L: -1.1, D: -2.85 with 20° angle) were performed stereotaxically in WT and *15q dup* mice: 50 μL of 0.5% CTB solution dissolved in saline was injected using a glass micropipette. The mice were anesthetized 5 days post-CTB injection, with an intraperitoneal injection of an anesthetic combination comprising 0.3 mg/kg medetomidine, 4.0 mg/kg midazolam, and 5.0 mg/kg butorphanol. Then, they underwent cardiac perfusion fixation with 4% PFA in 0.01 M PBS. The post-fixed brains were rinsed with saline and cryoprotected overnight with 30% sucrose in PBS at 4°C. Brain sections (50 μm) were cut using a sliding microtome (HM430; Thermo Fisher Scientific, Waltham, MA, USA). Immunostaining was performed as previously described.

### Statistical analyses

Statistical analyses were performed using PRISM 9 software (GraphPad Software, La Jolla, CA, USA). Data are expressed as the mean ± standard error of the mean. For comparisons, Student’s *t*-test, one-way analysis of variance (ANOVA) followed by Tukey’s post-hoc test, or two-way ANOVA followed by Bonferroni’s multiple comparison test were performed. P values < 0.05 were regarded as statistically significant.

## Data availability

The data generated in this study are available in the article, Supplementary Information and Source data. Source data are provided with this paper.

## Acknowledgments

We gratefully acknowledge the work of the present members of our laboratory, Dr. Shigefumi Yokota and Dr. Kenichiro Kuwako. We also acknowledge the technical expertise of the Interdisciplinary Center for Science Research, Head Office for Research and Academic Information, Shimane University.

## Authors’ contributions

Y.O. conceived and performed the experiments, analyzed the data, and wrote the paper with input from all authors. K.K. and R.K performed electrophysiological analyses. X.L performed the immunostaining and data analysis. T.T. interpreted results and wrote the manuscript. M.F. conceived, designed the project, secured funding, and wrote the manuscript.

