## Supplementary figures and images for "Aberrant axon initial segment is an indicator for task-independent neural activity in autism model mice"

### supplemental Figure1 20240215.tif

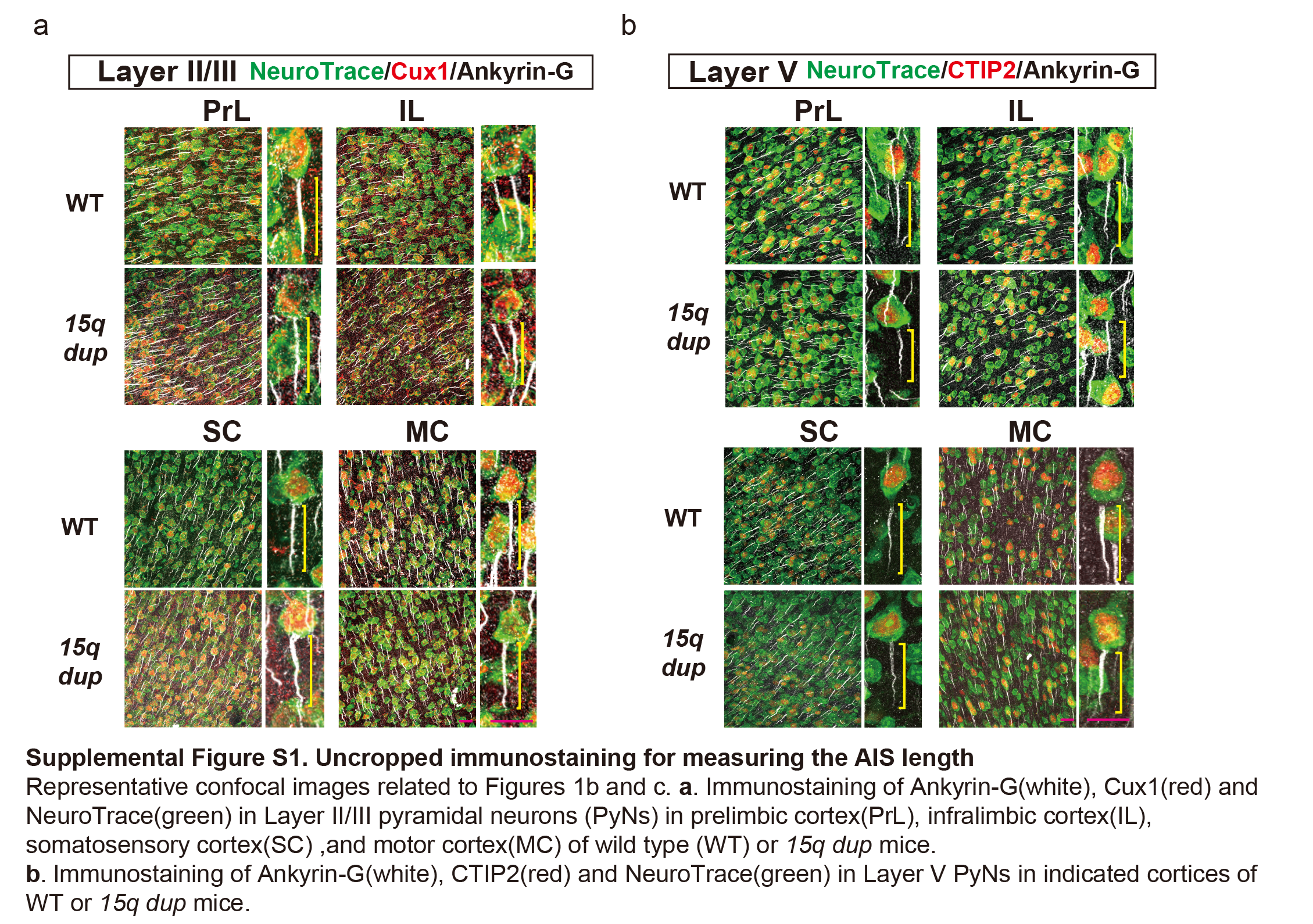

### supplemental Figure2 20240215.tif

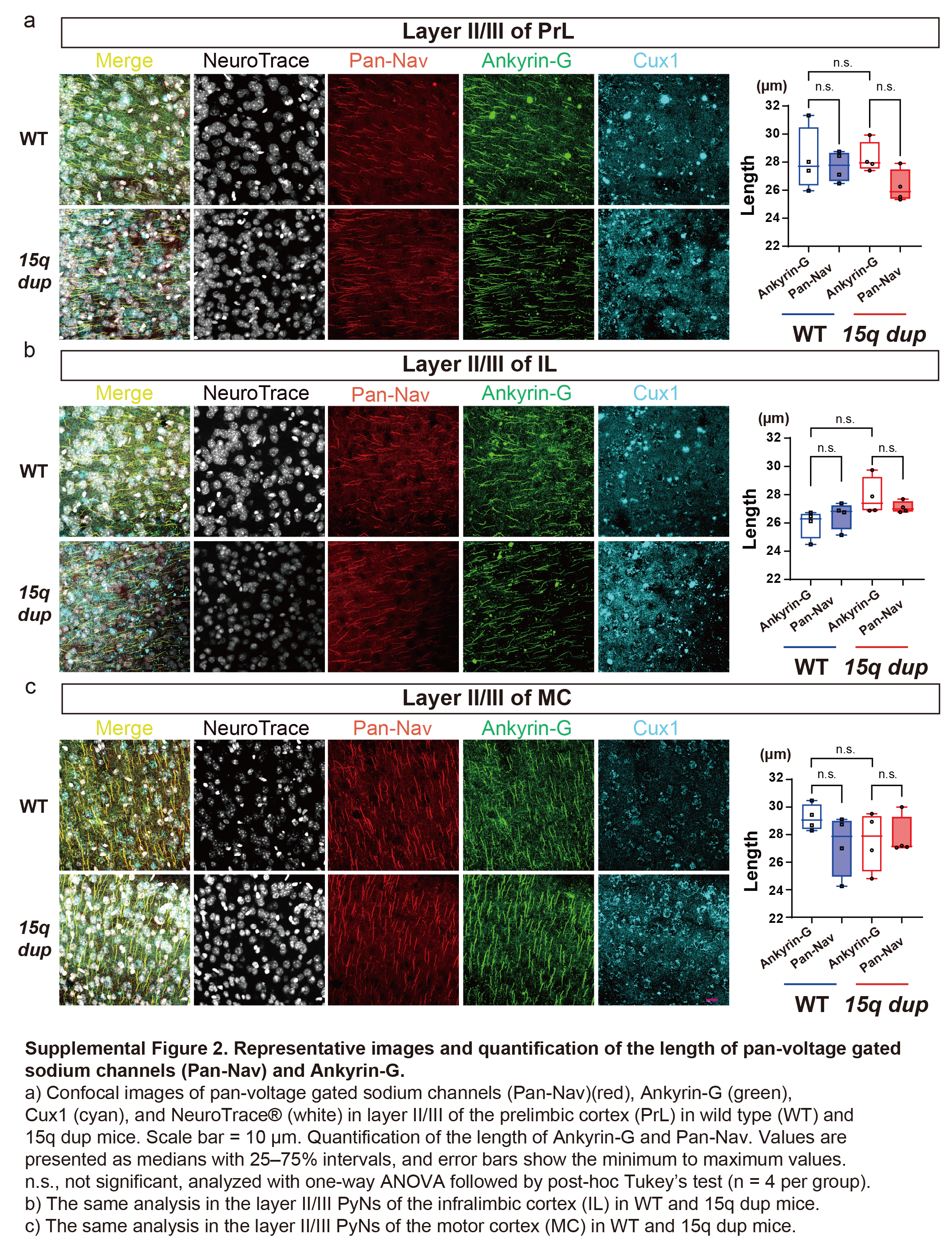

### supplemental Figure3 20240215.tif

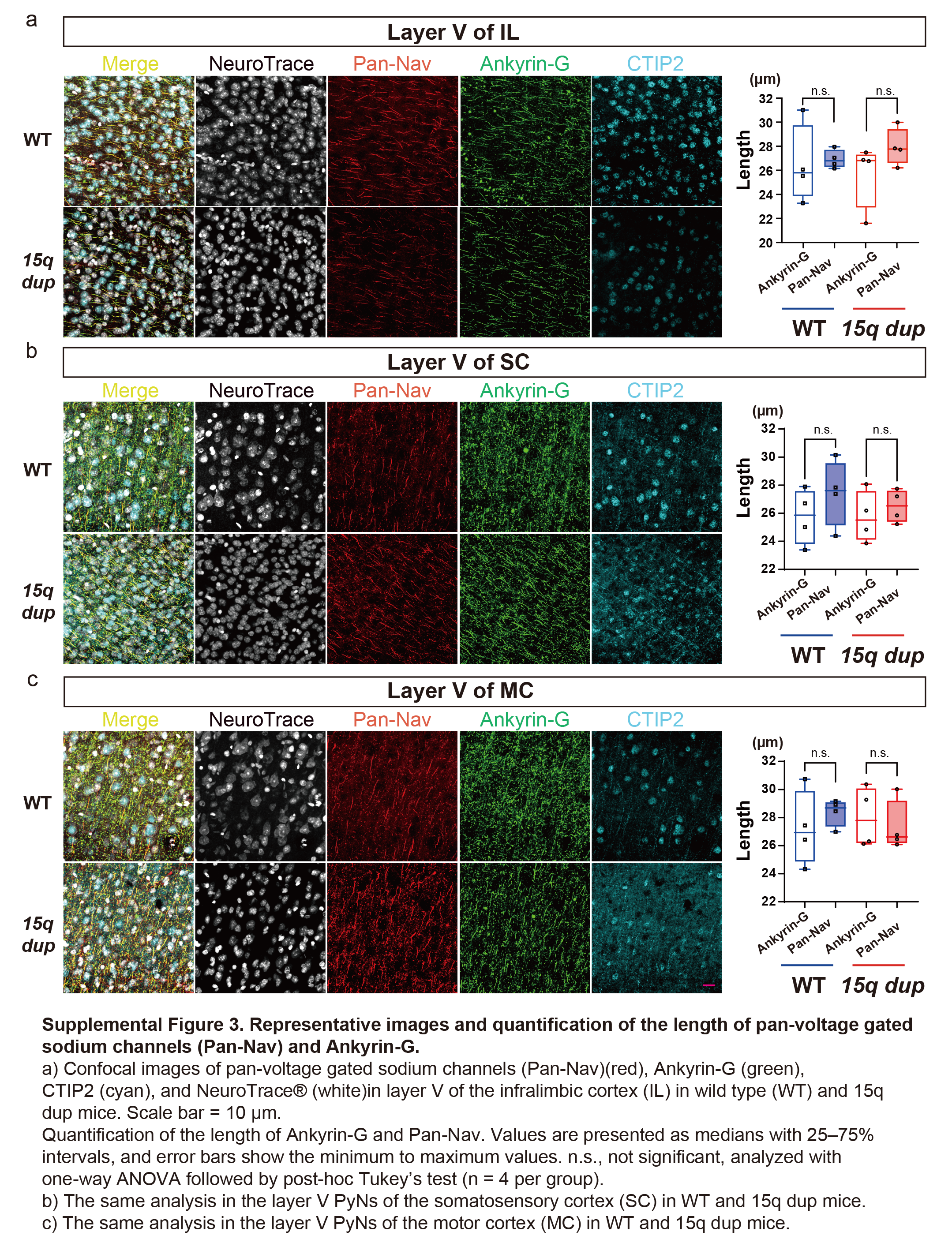

### supplemental Figure4 20240215.tif

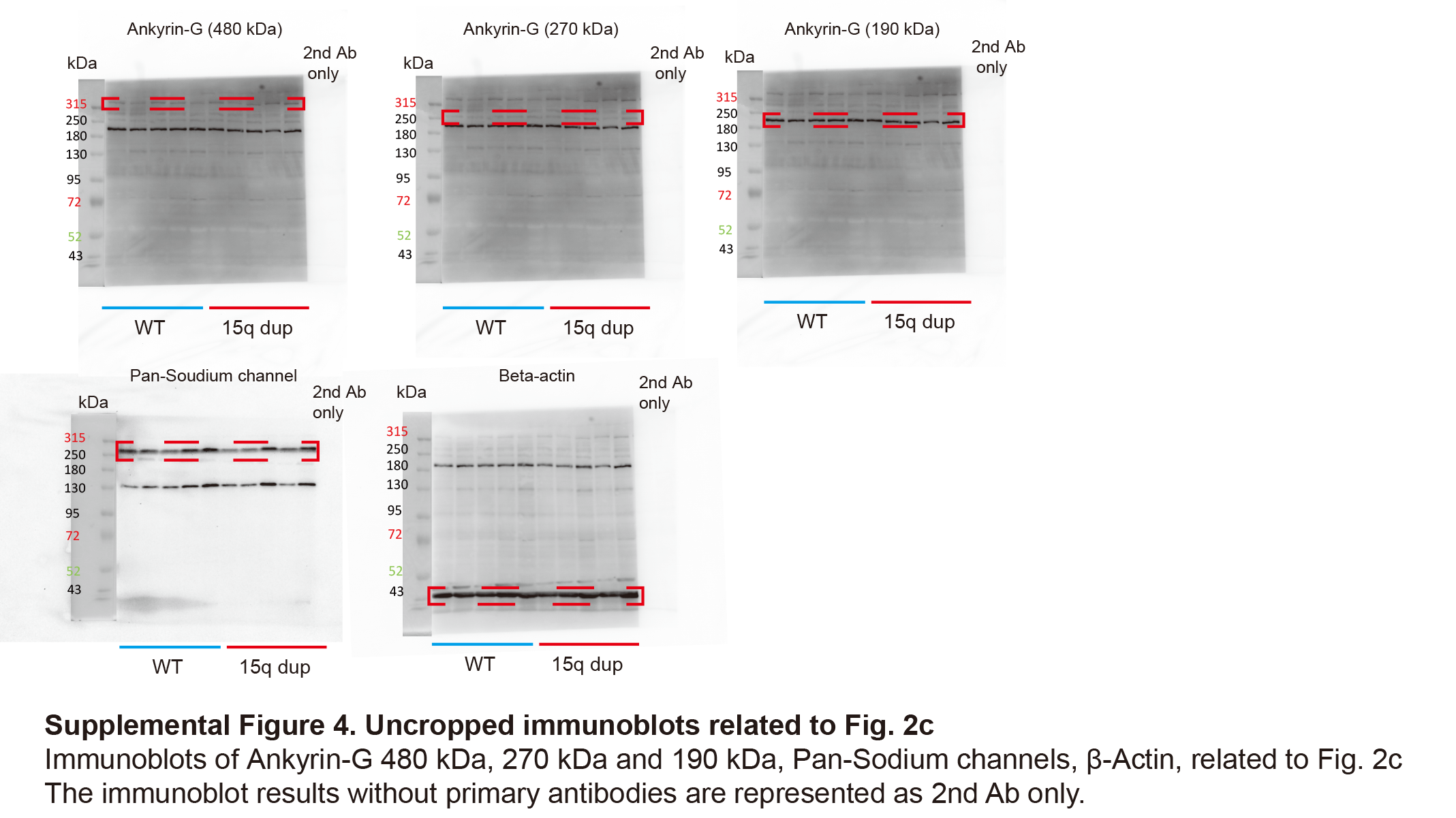
